# Metabolic response of microglia to amyloid deposition during Alzheimer’s disease progression in a mouse model

**DOI:** 10.1101/2023.05.12.540407

**Authors:** Kaitlyn M. Marino, Jayne M. Squirrell, Jenu V. Chacko, Jyoti W. Watters, Kevin W. Eliceiri, Tyler K. Ulland

## Abstract

Alzheimer’s disease (AD) drives metabolic changes in the central nervous system (CNS). In AD microglia are activated and proliferate in response to amyloid β plaques. To further characterize the metabolic changes in microglia associated with plaque deposition *in situ*, we examined cortical tissue from 2, 4, and 8-month-old wild type and 5XFAD mice, a mouse model of plaque deposition. 5XFAD mice exhibited progressive microgliosis and plaque deposition as well as changes in microglial morphology and neuronal dystrophy. Multiphoton-based fluorescent lifetime imaging microscopy (FLIM) metabolic measurements showed that older mice had an increased amount of free NAD(P)H, indicative of a shift towards glycolysis. Interestingly in 5XFAD mice, we also found an abundant previously undescribed third fluorescence component that suggests an alternate NAD(P)H binding partner associated with pathology. This work demonstrates that FLIM in combination with other quantitative imaging methods, is a promising label-free tool for understanding the mechanisms of AD pathology.

## Introduction

Alzheimer’s disease (AD) is a neurodegenerative disease that leads to systemic metabolic dysfunction1. AD is thought to be primarily driven by the accumulation of extracellular amyloid beta (Aβ) plaques occurring during the prodromal phase of disease, followed by the formation of intraneuronal neurofibrillary tau tangles (NFTs) at or around the time cognitive decline begins. Combined, these pathologies are thought to result in neuronal loss and cognitive decline. Additionally, AD is characterized by the reduction of global glucose metabolism resulting in reduced glycolytic flux, as confirmed by histology and fructose-deoxyglucose positron emission tomography (FDG-PET) 2–4. Although these studies identify changes in systemic and global brain glucose metabolism in the context of AD, the specifics of what cell types contribute and when during disease progression they occur remains ambiguous. Neurons are dystrophic within the AD brain, so we looked to examine what other cell types could contribute to this dysfunction, specifically glia.

Microglia are long lived, resident innate immune cells of the brain. During development they prune excess synapses 5 while under homeostatic conditions they continuously surveil the brain, containing foreign material and pathogens ^6,7^. Microglia have critical and diverse functions throughout the lifespan and have been implicated in several diseases, as their activation results in inflammation and subsequent changes within the microenvironment ^8–11^. While the role of microglia in neuro-pathologies is not fully understood, advanced sequencing has established microglia as a heterogeneous population in disease states, including AD and Amyotrophic Lateral Sclerosis ^12–17^. As microglia adopt different activation states their functions, such as cytokine production and phagocytic capacity, are altered ^15,17^.

While microglial activation states in the AD brain evolve with disease, it is known that microglia can both phagocytose Aβ fibrils and promote inflammation via inflammasome activation in the plaque microenvironment ^18,19^. Systematic inflammation can disrupt microglia by reducing their phagocytic capacity; however, in states of acute stress, microglia can switch their primary metabolism from oxidative phosphorylation to glycolysis, generating a surge of ATP production ^20–22^. In mouse models of AD that mimic Aβ deposition, microglia increase glucose utilization and downregulate homeostatic genes, like sall1 ^15,23,24^. Further, genes associated with microglial activation and microglial induced inflammation, such as IL-1β and mTOR, correlate with increased glycolytic metabolism ^25^.

Critically, metabolic dysfunction is implicated in a variety of pathologies. While the specifics may differ, ultimately, this dysfunction results in alterations to ATP production via major metabolic pathways *i*.*e*. glycolysis, the tricarboxylic acid (TCA) cycle, oxidative phosphorylation, and the electron transport chain. While systemic differences in the usage of these pathways can be readily detected via neuroimaging, identifying changes in the utilization of major metabolic pathways at a single cell level, within intact tissue, during disease development is more challenging. Single cell sequencing technologies have looked to address this but are limited in spatial and temporal resolution. Fluorescence lifetime imaging microscopy (FLIM) is a label-free multiphoton-based technology which can be used to examine cellular metabolism by visualizing the endogenous fluorescence of the ubiquitous cofactor nicotinamide adenine dinucleotide phosphate (NAD(P)H). FLIM analyses can evaluate the relative ratio of free (NAD(P)H or reduced NAD^+^) and bound NAD(P)H within live or fixed samples, which is indicative of the primary metabolic state *i*.*e*. the levels of glycolytic or oxidative metabolism used in the cell ^26,26– 28^. FLIM has been used previously in both *in vitro* and *in vivo* experiments to elucidate changes in NAD(P)H metabolism in several disease contexts, including the examination of subcellular metabolism in neurons and glia ^29–31^. To uncover microglia specific metabolic alterations within AD pathology, we utilized both confocal microscopy and FLIM in cortical tissue from 5XFAD mice, which models rapid amyloid plaque deposition. The result is a spatial and temporal look at the metabolic differences in the microglial response to Aβ plaques. We characterize microgliosis with increased plaque burden and demonstrate that microglial activation generates a unique signature, which does not correspond to oxidative phosphorylation or glycolysis.

## Methods

### RESOURCE AVAILABILITY

#### Lead contact

Further information and requests for resources and reagents should be directed to and fulfilled by the lead contacts; Kevin Eliceiri and Tyler Ulland

#### Materials availability

This study did not generate unique reagents.

#### Data and Code availability

The raw microscopy data and any additional information required to reanalyze the data reported in this paper is available from the lead contacts Kevin Eliceiri and Tyler Ulland upon request.

### EXPERIMENTAL MODEL DETAILS

#### Animals

WT mice (C57/Bl6; JAX Stock #000664) were crossed with 5XFAD mice ^32^ obtained from the Mutant Mouse Resource and Research Center at JAX (B6SJL-Tg(APPSwFlLon,PSEN1*M146L*L286V)6799Vas/Mmjax; Stock #034840-JAX) and maintained at the Biomedical Research Model Services (BRMS) at the University of Wisconsin. Age and sex matched (male and female) littermate controls were used for all experiments. All animals were housed according to the University of Wisconsin Institutional Care and Use Committee guidelines under the approved protocols V005173 and M006091. Mice were maintained on a 12-hour light/12-hour dark cycle with food and water provided *ad libitum*.

## METHOD DETAILS

### Tissue Preparation and Immunohistochemistry

Mice were anesthetized with isoflurane and perfused with cold phosphate buffered saline (PBS) with 1 U/ml heparin (Hospira Pharmaceuticals; NDC 0409-2720-32). As reported previously (Shippy et al., 2020), brains were fixed in 4% paraformaldehyde for 48 hours, rinsed with PBS, sunk in 30% sucrose and frozen in a 2:1 30% sucrose/Tissue Tek OCT compound (Sakura #4583) solution. Brains were sectioned to 40 µm (Leica) and stored in cryoprotection buffer (30% ethylene glycol, 30% sucrose in PBS). Sections were blocked (3% BSA, 0.15% Triton-X100 in PBS) for one hour and incubated with primary antibody (anti-Iba1(Abcam ab5076; 1:1750) or anti-BACE-1 (Abcam 108394, 1:500)) overnight at 4°C. Sections were rinsed with PBS and incubated with secondary antibody (Alexa Fluor donkey anti-goat 568 (Thermo Fisher A-11057; 1:750) or DyLight 488 Goat anti-Rabbit (Vector Laboratories DI-1488, 1:1000)) for 1 hour. Sections for confocal microscopy were incubated with MethOxy X04 (Tocris Biological #4920; 1:1000) and TO-PRO-3 (Thermo Fisher #T3605; 1:1000) during the same interval. Sections were rinsed with PBS and mounted with Fluoromount G (SouthernBiotech, Cat. No. 0100-001) and #1.5 coverslips (Fisherbrand, Cat. No 12544B).

### Confocal Microscopy

Confocal images were taken on the Nikon A1R confocal microscope in the University of Wisconsin Optical Imaging Core (Madison, WI). Cortical images were taken using the 20X air 0.75 NA or 60X oil 1.4 NA objectives as Z stacks with a step of 0.7 µm and exported to Imaris (Bitplane 9.5.1) or Image J ^33^ for analysis.

### Cluster Analysis

As described previously ^34^ z-stack images (20X; 2.42 pixels per µm were analyzed in Imaris. The nuclei (TO-PRO-3) and microglia (Iba-1) channels were colocalized; a 3D cell count was generated using the spots function. For plaque analyses, the surfaces tool was used in the methoxy X04 stain channel to generate plaque surfaces. All plaque statistics (sphericity, area, quantity) were recorded for each image.

### Morphology Analysis

Several high magnification (60X; 9.67 pixels per µm) confocal images from within the cortex were imported into Image J ^33^ using the Bio-Formats plugin ^35^. Using the SNT plugin ^36,37^ individual microglia were traced in 3-dimensions. Using the batch analysis feature, single cell measurements (number of branch points and number of branches) and Sholl analyses (3 µm radius) was generated.

### BACE-1 Analysis

High magnification (60X; 7.25 pixels per µm) confocal images from within the cortex were taken, ensuring that plaques were well within the imaging field. In Imaris, BACE-1 staining was rendered as a surface (0.5 µm detail) and quantified. For individual plaque analyses, the methoxy X04 channel was used to make a surface of a plaque and was dilated by 15 µm. The BACE-1 volume was colocalized with the expanded plaque. This overlapping volume was normalized to the size of the dilated plaque and reported as a percentage.

### FLIM Imaging

Single focal plane images were obtained on a custom-built multiphoton microscope equipped with a Ti:Sapphire laser (Chameleon UltraII, Coherent Inc, Santa Clara, CA), described previously ^38^, using a 40X water immersion lens (1.2 NA, Nikon, Melville NY). Iba-1 fluorescence was imaged using multiphoton excitation at 800nm and collected using a narrow-band emission filter centered at 615 (Semrock, FF01-615/20-25). The corresponding NAD(P)H lifetime was captured with an excitation of 740nm and emission filtering through a 457nm narrow band emission filter (Semrock, FF01-450/70-25) over a collection interval of 60s in the same region of interest. The laser is scanned on the sample using galvanometric scanners coupled to lab-made acquisition software ‘WiscScan,’ and the emission signal is collected using a high gain photon counting GaAsP PMT (Hamamatsu, H7422P-40). The arrival time of each photon was calculated using Becker Hickl TCSPC electronics (SPC-150 board). Image standards were taken at each imaging session to minimize instrument variability. Images from a known fluorescence target (Chroma slide/1.1 ns) and a second harmonic generation emitter (Urea crystals, 0 ns) were used as lifetime standards. Multiple sets of images were collected in approximately the same location in the cortex using the midline as a reference point. 3 images at least 200 microns apart were collected for each section and 2 sections were used for each animal.

### FLIM Analysis

The FLIM data (x,y,T) data from NAD(P)H channel was analyzed using SPCImage 8.5 (Becker & Hickl GmBH, Berlin) and transformed into phasor coordinates (x,y,g,s). The images were saved in a spreadsheet after being mapped to their matching antibody immunofluorescence image Ab(x,y)->FLIM(x,y) (as a json file). The animal labels, region labels and image labels were preserved and accessible using a unique image ID. Each image’s FLIM data, which consists of 256×256 XY-pixels, was downsampled to 64×64 before being compared to the matching antibody-stained image. Fluorescence image from Iba-1+ cells from the same field of view with an intensity threshold (20%) was used as a mask for microglia regions. Additional steps to remove faulty pixels from FLIM data and an evaluation of image-landmark overlap was performed for the dataset manually. For statistical analyses and plotting, the combined array of both images (two channels NAD(P)H/Antibody stain) was saved as [2 × 1024] per image (field of view). Additional parameters (mean lifetime, phasor coordinates, fit results, goodness of fit and others) were added as columns to the pixels as [n x 1024] in the data frame, where n is the number of parameters. The phasor coordinates from each (downsampled) pixel mapped to the universal phasor circle was studied in three steps. The phasor distribution aligned across the free-NAD(P)H: bound-NADH(P)H line (0.4 ns, 2.95 ns) with a cohort of points pulling away from the line, and three component phasor analysis was required. We verified this was not from background by applying the microglia mask from antibody images. For the three-component analysis; first, a triangle encompassing all the phasor points was drawn; next, the 2D phasor distribution was fitted to a gaussian mixture model using the vertices as initial centers; and finally, the orthogonal distance projection between each phasor point and the connecting line joining the free-NAD(P)H and bound-NAD(P)H lifetime measured the two-component estimate of NAD(P)H fraction. The mixture model fit identified three means ([0.49, 0.39], [0.54, 0.39], [0.55, 0.37]). This phasor vector math to identify multiple clusters is detailed in Vallmitjana et al., 2021. Three major outputs are produced by this process for this three-species phasor analysis: a) the extrapolated free: bound fraction of NAD(P)H without the influence of the third cluster; b) the strength of the third cluster (fraction of pixels in the cluster, based on the distance from the third cluster); c) set of images that maps the phasor location, cluster intensity, and other FLIM parameters at each pixel.

### Microglia microarray

Microglia microarray data was accessed from GEO (GSE65067), the subsets of WT and 5XFAD microglia were compared ^40,41^. Subsequent analyses compared specific genes relating to glycolysis, the TCA cycle and oxidative phosphorylation identified in the literature between WT and 5XFAD microglia with a false discovery rate (FDR)<0.05 ^40,42–44^.

## QUANTIFICATION AND STATISTICAL ANALYSIS

### Statistics

Graph Pad Prism (version 9.4.1) was used for statistical analyses (unpaired *t-*test, one-way or two-way ANOVA with Tukey multiple comparisons) at a significance of *p*<0.05. Python (Python Software Foundation) was used for Phasor analysis with help of FlimLib library ^45^ for FLIM analysis, Pandas ^46^ for database management and statsmodel library ^47^ for phasor-related statistics outputs. Comparison groups included male and female mice.

## KEY RESOURCE TABLE

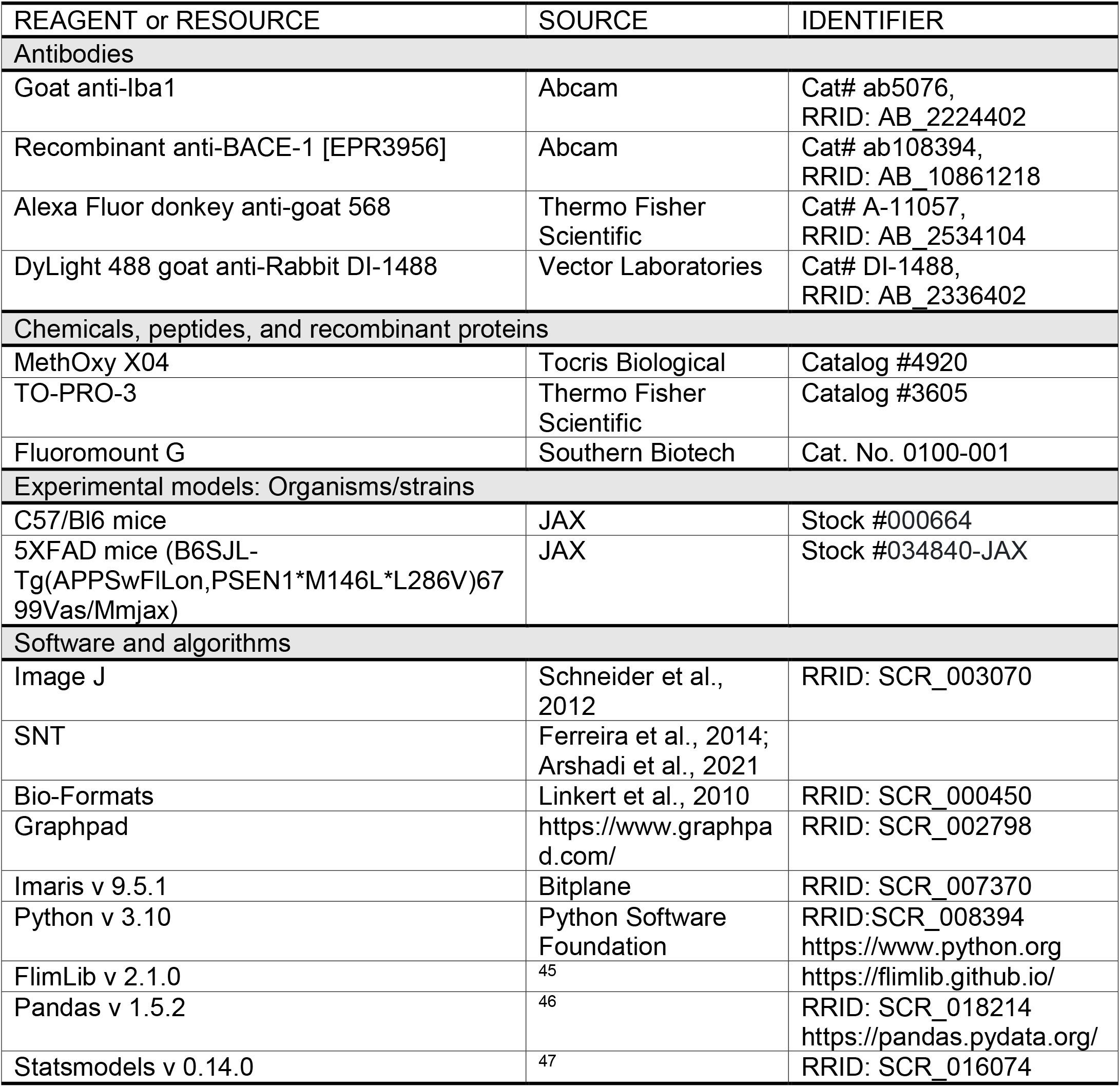

## Results

### Aβ plaque load and density increase with age

To document the progression of amyloid deposition over time, the number of MethOxy X04-labeled plaques (Figure 1A) within a high magnification field (422×422 µm) was counted in Imaris (Bitplane 9.5.1). The number of plaques per field increased at each age (2-,4- and 8-months; Figure 1B; One-way ANOVA, *p*<0.0001). Since no significant differences were indicated by sex, all further analyses include both sexes. Furthermore, using the surface rendering tool in Imaris to assess plaque volume, we found that plaque volume significantly increased between 2 and 4 months of age, but did not increase from 4 and 8 months (Figure 1C; One-way ANOVA, *p*<0.05). The sphericity of the plaques did not change (Figure 1D; One-way ANOVA, p>0.05), so the plaque morphology itself was not changing, just the size and quantity. Since the plaque number increased between 4 and 8 months but plaque volume did not, we measured the global amyloid burden using large scale imaging of the cortex of 4- and 8-month animals (Figure 1E-F). Few or no plaques were seen in 2 months animals (Figure 1A, B) so these were not included in this analysisA significantly higher percentage of the cortex contained plaques in 8-month 5XFAD compared to 4-month 5XFAD mice (Figure 1F; Unpaired *t*-test, *p*<0.01). These results characterize the rate of plaque accumulation within the ages of 5XFAD mice used in the analyses below.

**Figure 1.**
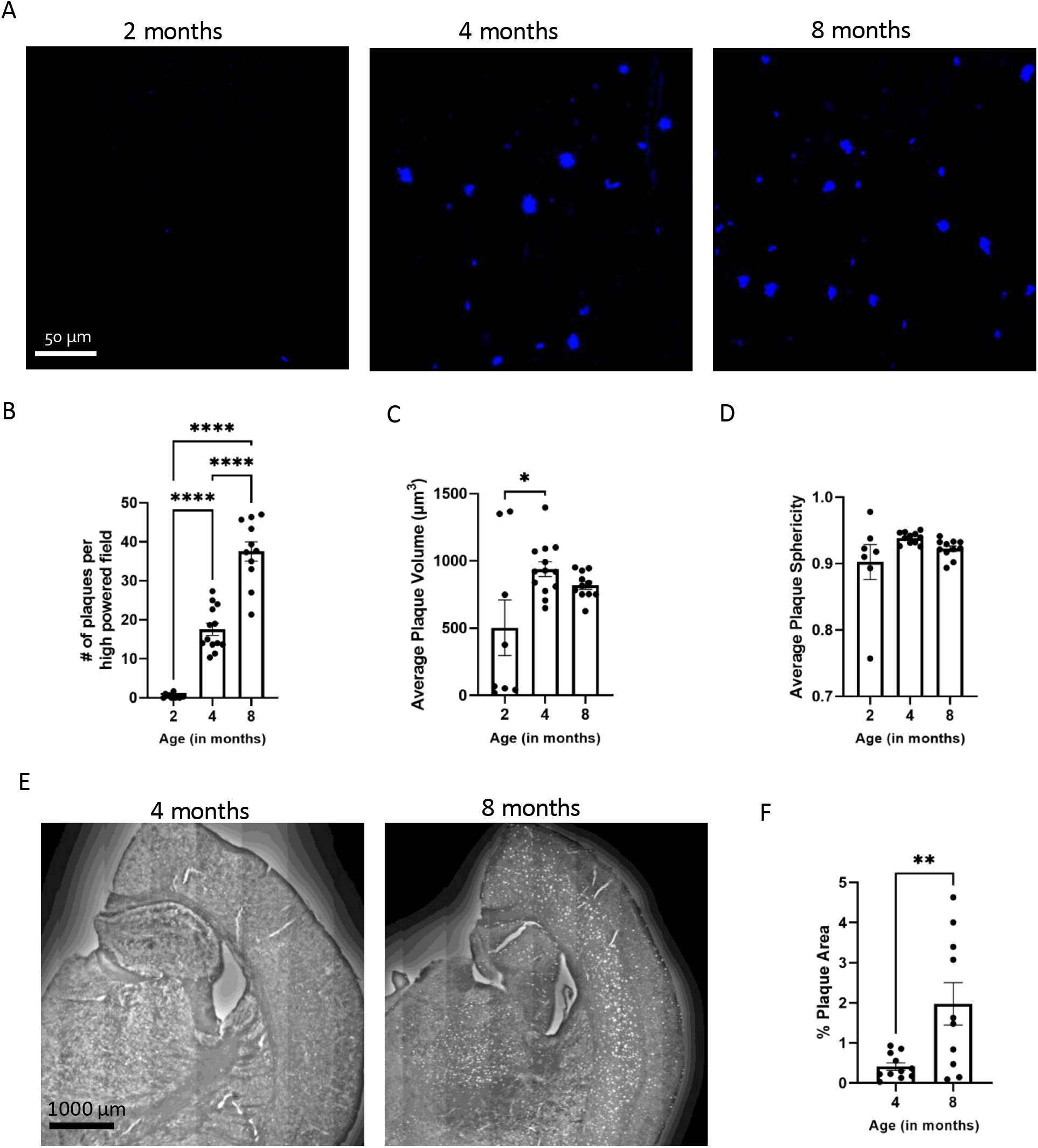
Plaque quantity and characteristics change with age. (A) Confocal images of methoxy-X04 staining (blue) in a high-power field in the cortex of 5XFAD mice at 2 months, 4 months, 8 months of age. Imaris quantification of plaque number per field (B), average plaque volume (C) and plaque sphericity (D). Large scale images of the 4-month and 8-month 5XFAD cortex (E), and quantification of the proportion of the cortex that contains plaques (F). Each point represents at least 3 images from a single animal (B-D). All data is presented ± SEM. One-way ANOVA with Tukey multiple comparisons (B-D), unpaired t-test (F), ****p<.001, **p<0.01, *p<0.05.

### Plaque deposition results in microgliosis and decreased morphological complexity

Microglia respond to amyloid deposition and are known to have many roles, so we characterized their response with the progressively increasing pathology documented in Figure 1. The number of Iba-1^+^ cells (microglia) within a high-magnification field as well as their proximity to a plaque were quantified in Imaris. The number of microglia in the cortex of 5XFAD animals increased compared to WT animals at 2-, 4-, and 8-month of age, respectively (Figure 2 A-B; ANOVA, *p*<0.05). To assess the interactions between microglia and plaques, we performed a cluster analysis to determine the number of microglia within 15 µm of a Aβ plaque. However, due to the sparse number of plaques in the 2-month cortices, the interactions between microglia and plaque were only characterized in 4- and 8-month-old animals. The average number of microglia within 15 µm of each plaque in the imaging field and the total number of microglia within 15 µm of the plaque surface increased at 8 months of age (Figure 2C-D, *t*-test, *p*<0.0001). However, the number of microglia greater than 30 µm of a plaque surface, a radius that has been shown to contain microglia unaffected by plaque pathology ^48^, remained unchanged between 4 and 8 months (Figure S2). Therefore, the microglia response to Aβ increases with age (Figure 2C-D, unpaired *t*-test, *p*>0.05) but the number of non-plaque associated (>30µm) microglia does not change (Figure S2).

**Figure 2.**
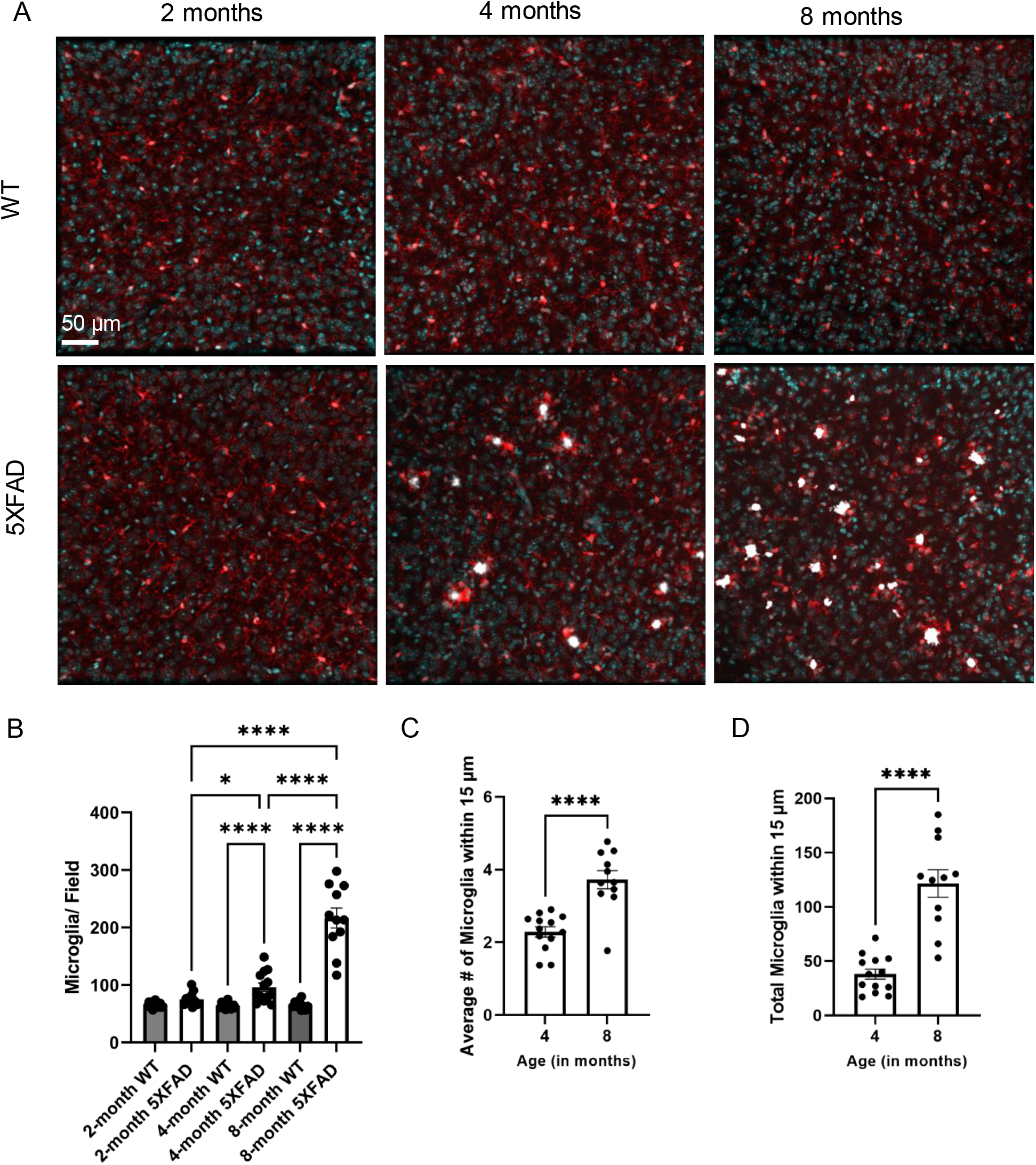
Microglial population and response to Aβ deposition increases with age. Confocal imaging of the cortex in 2, 4 and 8-month animals using Iba-1 (red), methoxy X04 (white) and TO-PRO-3 (blue) (A). Comparisons of microglia count from per imaging field at 2-, 4-, and 8-month ages (B). Cluster analysis of microglia around a plaque. Calculated average number microglia (C) and the total number of microglia (D) within 15 µm of a plaque. All data is presented ± SEM and each point represents 3 images. ANOVA with Tukey correction; Unpaired t-test, ****p<0.0001, ***p<0.001, **p<0.01, *p<0.05.

To characterize individual microglia characteristics, we analyzed cell morphology (Figure 3). High magnification images captured 1-2 cells in the cortex; for 5XFAD mice, only cells within 30 µm of a plaque surface were selected while WT microglia were randomly selected. Microglia traces were generated for microglia confirmed with nuclei (TO-PRO-3) using the SNT plugin. Using SNT, cells were compared for complexity via branch point quantification and using a 3-dimensional Sholl analysis ^37^, which quantifies the number of intersections at concentric radii. These data were plotted for each cell and the area under that curve was reported. Analyses indicate that microglia from WT mice were significantly more complex than plaque associated microglia in 5XFAD mice at 4- and 8-months of age (Figure 3B-C; ANOVA).

**Figure 3.**
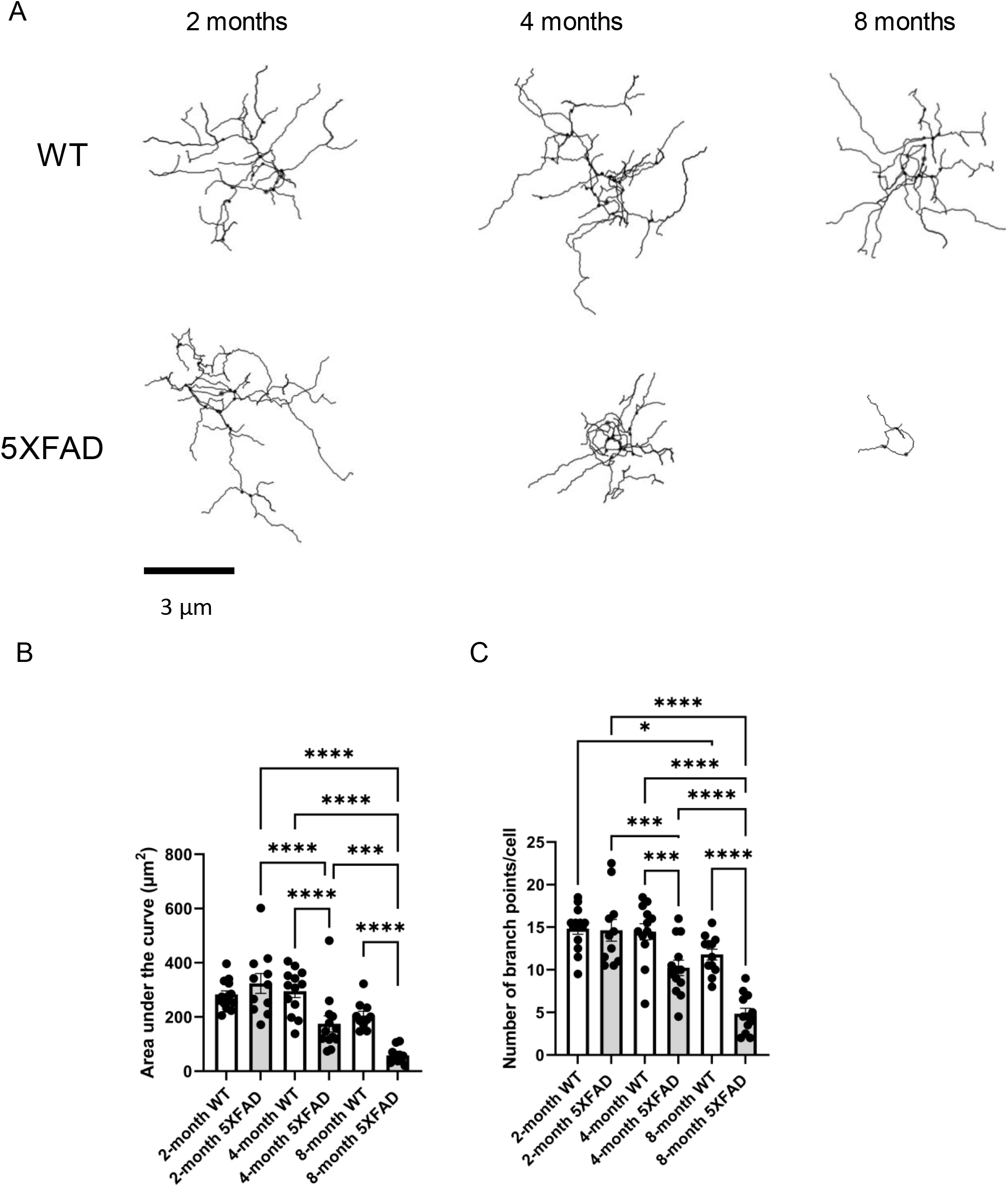
5XFAD microglia lose complexity with age. Representative WT and 5XFAD cortical microglia at 2-,4-, and 8-months of age, created using SNT (A). Area under the curve from a Sholl analysis (B). Total number of branch points per cell (C). All data is presented ± SEM and each point represents 2 microglia. ANOVA with Tukey corrections, ****p<0.0001, ***p<0.001, **p<0.01. *p<0.05.

### Neuronal health is differentially impaired with amyloid deposition

To evaluate how disease progression impacted neuronal health in the 5XFAD model, BACE-1 was used to quantify dystrophic neurites near plaques. BACE-1 labels inactive, presynaptic terminals found within the plaque microenvironment ^49,50^. BACE-1 localizes in the peri-plaque region ^49,51^; we focused initial analyses on the BACE-1 volume within 15 µm from the surfaces of individual plaques. In Imaris, 3D surfaces were created from the methoxy X04 and BACE-1 channels, respectively. The percentage of the volume occupied by BACE-1^+^ pixels within 15 µm of a plaque surface was calculated. The volume of BACE-1^+^ pixels within 15 µm of a plaque was decreased in the cortex of 8-month 5XFAD animals compared to 4-month animals (Figure 4B, Unpaired *t-*test, *p*<0.05) consistent with previously published results ^52^. However, when global neuronal dystrophy was surveyed and the total volume of BACE-1^+^ pixels was quantified in the 5XFAD cortex, 8-month animals had significantly more BACE-1 positivity (Figure 4D, Unpaired *t*-test, *p*<0.001). Thus, increased global, but not local, neuronal dystrophy is seen in the 8-month cortex compared to 4-month by BACE-1 staining, consistent with other markers of dystrophy in the 5XFAD mouse model ^52^.

**Figure 4.**
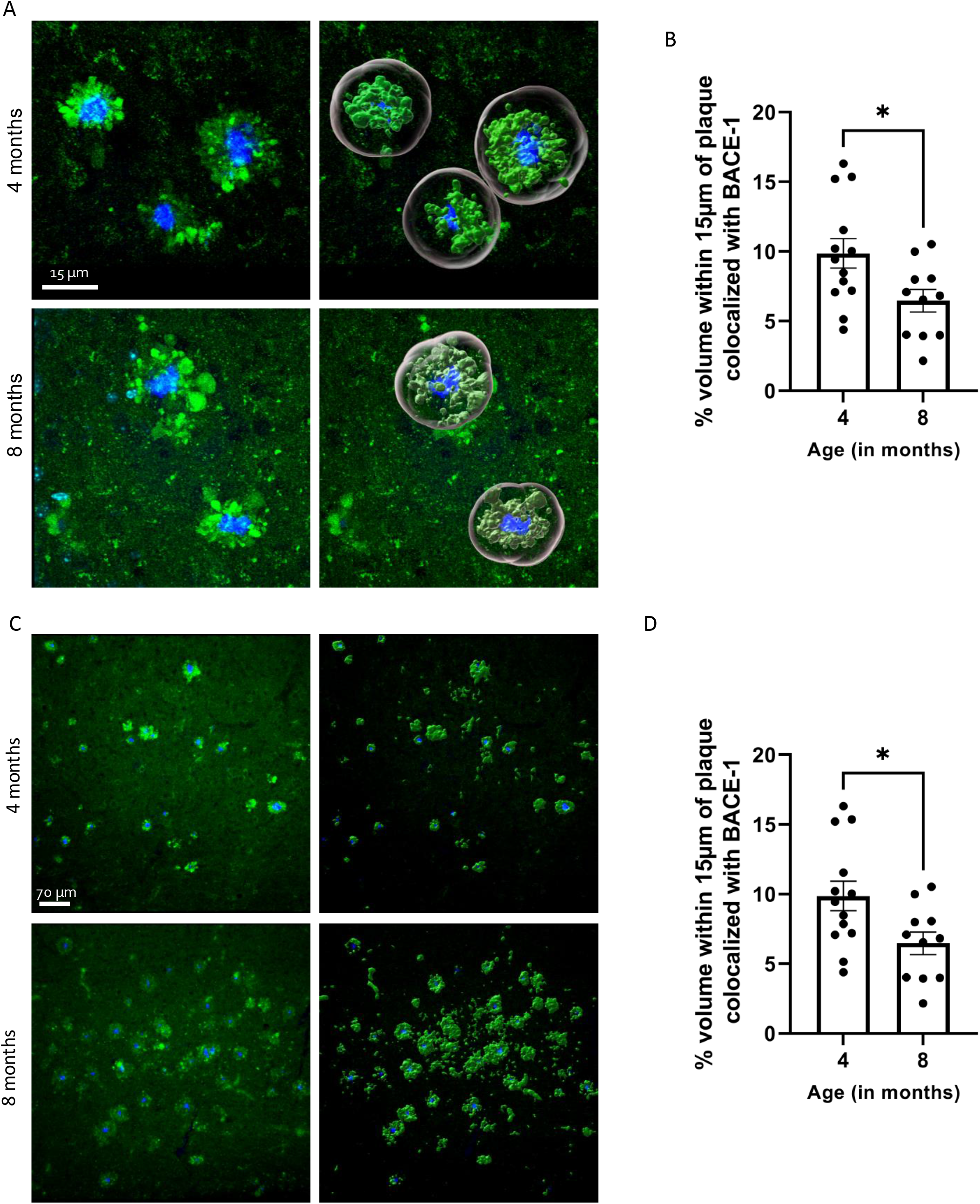
Global increase in dystrophic neurite quantity in 8-month cortex. Confocal images of dystrophic neurites (BACE-1; green) surrounding plaques (Methoxy-X04; blue) in the cortex of 4- and 8-month 5XFAD mice (A). Imaris quantification of the percentage of the volume within 15 µm of a plaque that is occupied by BACE-1 stain (B). Each point represents >6 plaques. Confocal imaging of a lower power field (C). Quantification of the total volume of the field that is BACE-1^+^. Each point represents 2 images (D). All data is presented ± SEM; unpaired t-test ***p<0.001 *p <0.05.

### Microglia NAD(P)H metabolism is age and genotype dependent

Briefly, Iba-1 intensity images were used to generate the threshold that was then applied to the lifetime data, Representational images for each group are presented in Figure 5A. To quantitatively evaluate the NAD(P)H metabolism Iba-1^+^ pixels were plotted on a universal circle to visualize similar metabolic species (Figure 5B). With a two-component estimate of the data, the relative free/bound NAD(P)H ratios are extrapolated. When quantified, significant differences in the percentage of free NAD(P)H within Iba-1+ pixels are found at 8-months of age between WT and 5XFAD microglia populations (Figure 5C, ANOVA, p<0.05). Further, within the microglia of WT mice, this percentage of free NAD(P)H sequentially increased (Figure 5C, ANOVA, p<0.05). However, in 5XFAD microglia population the relationship significantly increased between 2- and 4-months of age (ANOVA, p<0.0001), but not from 4- to 8-months of age, suggesting that plaque deposition and microgliosis are significant contributors (Figure 1 and 2 respectively).

**Figure 5.**
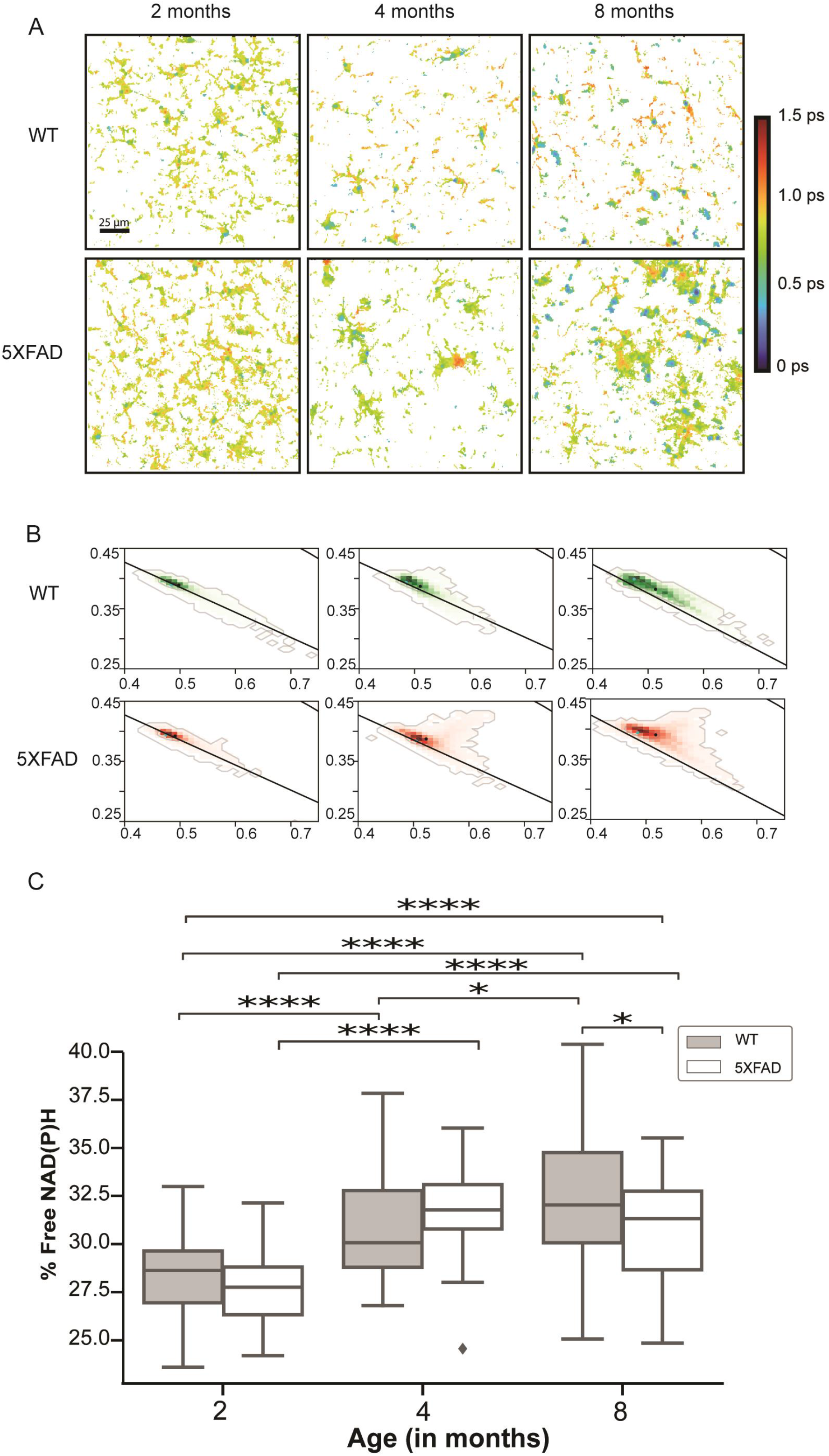
Altered NAD(P)H binding signature in aging microglia. Fluorescence lifetime within the Iba-1^+^ pixels of a cortex ROI for 2-, 4-, and 8-month WT and 5XFADs. Phasor plots of the corresponding to the lifetime of iba1^+^ pixels for 2-, 4-, and 8-month wildtype and 5XFAD cortex images (B). Calculated percent free NAD(P)H in microglia by age and genotype (C) ANOVA with Tukey comparisons; ****p<0.0001, ***p<0.001, **p<0.01, *p<0.05.

### A third NAD(P)H signature is more abundant in the later stages of plaque deposition

Surprisingly, we found that the free and bound NAD(P)H clusters alone do not account for all of the FLIM components observed. When the data was initially visualized on the normal circle (Figure 5B) there was an indication of another metabolic species found orthogonal and slightly above the center of the line joining the free-NAD(P)H and bound-NAD(P)H lifetime. Visually, the third cluster, identified by the distance from that free vs bound line, is clear when the free and bound clusters are labeled (Figure 6A). In Figure 6B, the spatial localization of the unknown cluster is visualized by the cluster strength for the unknown and primarily localizes in areas where you would expect to see microglia responding to a plaque. The cluster strength is a function of how many pixels in the image are attributed to the cluster. When the percentage of iba-1+ pixels that are a part of this third component is quantified, we see that the microglia from 4- and 8-month 5XFAD mice have the most compared to WT mice (Figure 6C, ANOVA, p<0.001). Microglia from 2-month WT or 5XFAD mice almost no abundance and differences between WT, but not 5XFAD microglia are reported between the 4- and 8-month groups (Figure 6C).

**Figure 6.**
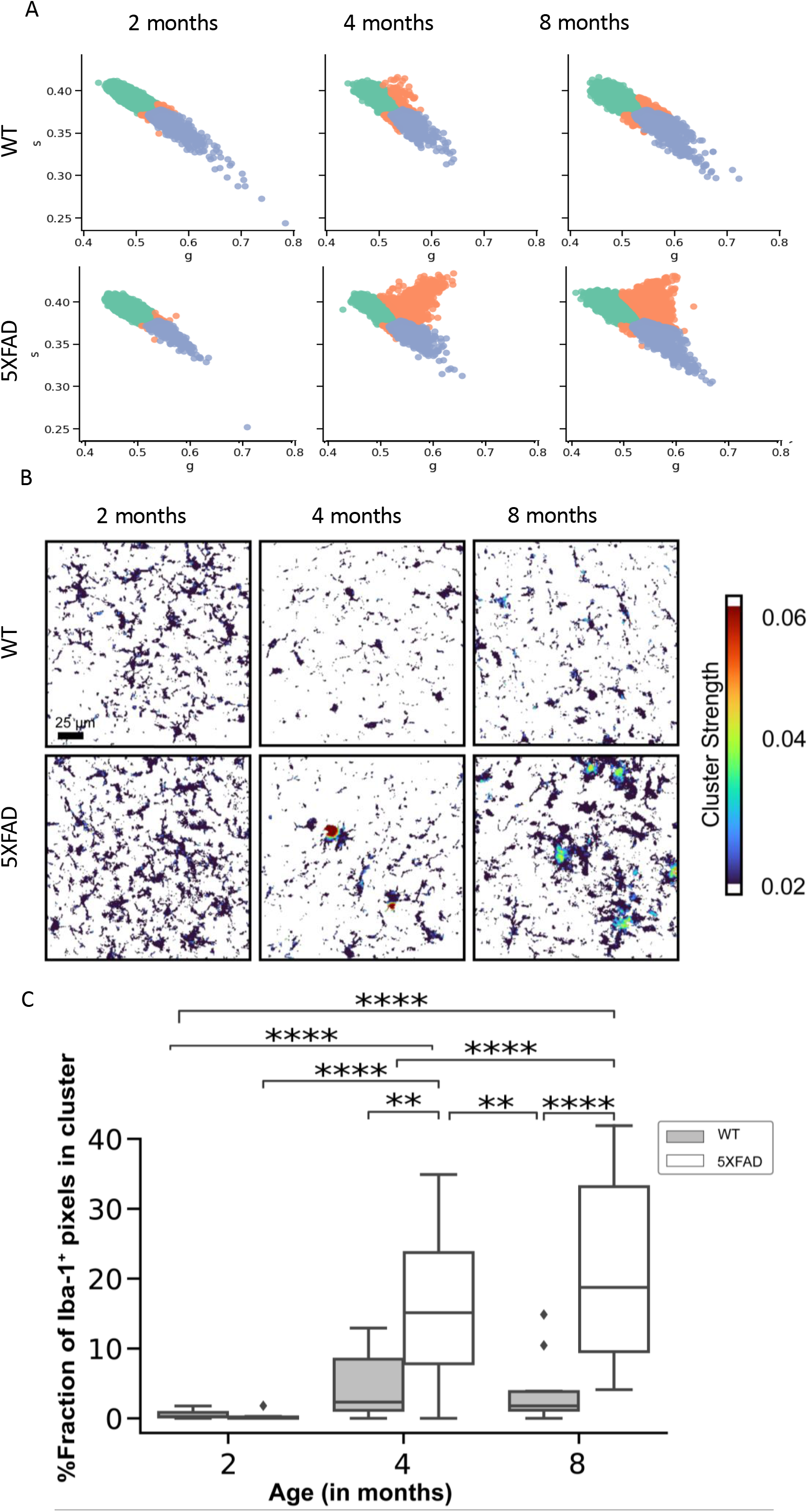
Clustering indicates a 3^rd^ component. Visual cluster distribution for Cluster 1 (green), 2 (purple) and 3 (orange; unknown) by age and genotype (A). Visual mask of the unknown cluster strength within the Iba-1^+^ area by age and genotype (B). Quantification of the percentage of the microglia pixels that contain the unknown cluster by age and genotype (C). ANOVA with Tukey comparisons; ****p<0.0001, ***p<0.001, **p<0.01, *p<0.05.

## Discussion

The 5XFAD model is well characterized (Forner et al., 2021; Oakley et al., 2006); however, the plaque deposition and microgliosis quantification presented in Figures 1-4 inform our FLIM analyses. Microglia quantity was increased compared to littermate controls throughout our time course and these microglia were more ameboid by morphologic analysis, supporting increased microglial activation. Further, the plaque burden in the cortex increased with age and the microglial response and global neuronal dystrophy increased with it. The analysis of microarray data to examine microglial metabolism (Figure S1) did not offer a clear target for classical imaging, therefore we turned to FLIM to further elucidate metabolic changes brought on by both aging and AD-like pathology.

Metabolic changes, including the Warburg effect, are well characterized in cancer cells and macrophages (Reviewed in Vander Heiden et al., 2009) and have been explored previously in microglia (Baik et al., 2019). We demonstrate the utility of FLIM to characterize the metabolic changes of microglia in relation to Aβ deposition in conjunction with traditional techniques. The relative increase of free NAD(P)H in wildtype microglia could support existing evidence that glucose utilization is reduced during the aging process, in part, as microglia become more glycolytic, perhaps as they use increasing quantities of fatty acid metabolism (Flowers et al., 2017; Goyal et al., 2017). However, we acknowledge that the animals in our analyses were relatively young, but the relative changes we captured with FLIM seem robust. The relative free NAD(P)H in microglia from 2- to 4-month 5XFAD animals is supported by previous evidence that that initially microglia use oxidative phosphorylation to address Aβ but shift towards glycolysis with increasing stress and inflammation to support proliferation (Baik et al., 2019; Cheng et al., 2021). Microglia are known to become tolerant to Aβ stress and this can cause increased oxidative phosphorylation utilization (Baik et al., 2019).

Critically, these data implicate a third, previously undescribed, component detectable in all cortical microglia, but at a significantly higher quantity in microglia from 4- and 8-month-old 5XFAD mice. While microglia in the AD brain are known to become more glycolytic (Baik et al., 2019), our data may indicate a previously overlooked, secondary NAD(P)H binding partner. This alternate signature is most abundant in 4- and 8-month 5XFAD microglia suggesting that it is driven primarily in response to increasing Aβ deposition. Further exploration of this third component/alternate binding partner would be of significant value in our understanding of amyloidosis. Unfortunately, this is currently challenging due to several factors. The use of additional fluorophores would distort or interfere with the NAD(P)H signal; any target could not be co-localized with a fluorescent microglia marker and since most targets would likely be ubiquitous within the brain and expressed in multiple cell types, the colocalization would be critical. As FLIM cannot distinguish between aerobic and anaerobic glycolysis, lactate metabolism could be considered; however, lactate metabolism is known to decrease in the presence of Aβ (Zhang et al., 2018). This does not appear to be the case, since this third component increases with amyloid burden.

While altered metabolism is found in the AD brain using other techniques, understanding the nuance of the changes at single cell resolution will enable better targets for improving AD pathology. The 5XFAD model is a rapid and robust model of amyloidosis and results in a strong glial response as we have demonstrated here. Microglia can have a wide variety of effects with pathology and uncovering those differences may lead to better identification and treatment of neuro-pathologies.

## Conclusions

Microglial metabolism changes with age and amyloid accumulation. By using FLIM, we are able to use little tissue to visualize single cell changes in high resolution while maintaining spatial and temporal control. These data offer a key insight into the microglial response to amyloid plaques and an opportunity to explore this further to better understand AD pathology.

## Acknowledgements

Special thanks to the UW ADRC for their support of this project and to Ellen Dobson for her assistance with the morphology analysis workflow.

## Authors’ contributions

KWE and TKU initiated the project concept and obtained research funding. KMM, JMS, JJW, KWE and TKU participated in study design discussions. KMM, JJW and TKU generated and collected samples. KMM preformed confocal microscopy imaging. JMS preformed the FLIM imaging experiments. KMM did all confocal microscopy analyses and statistics. JVC performed the FLIM phasor analysis and image statistics for the FLIM dataset. KMM wrote the manuscript, JMS, JVC JJW, KWE, and TKU participated in manuscript review and discussions.

