## Supplemental Figures for "Metabolic response of microglia to amyloid deposition during Alzheimer’s disease progression in a mouse model"

**S1**

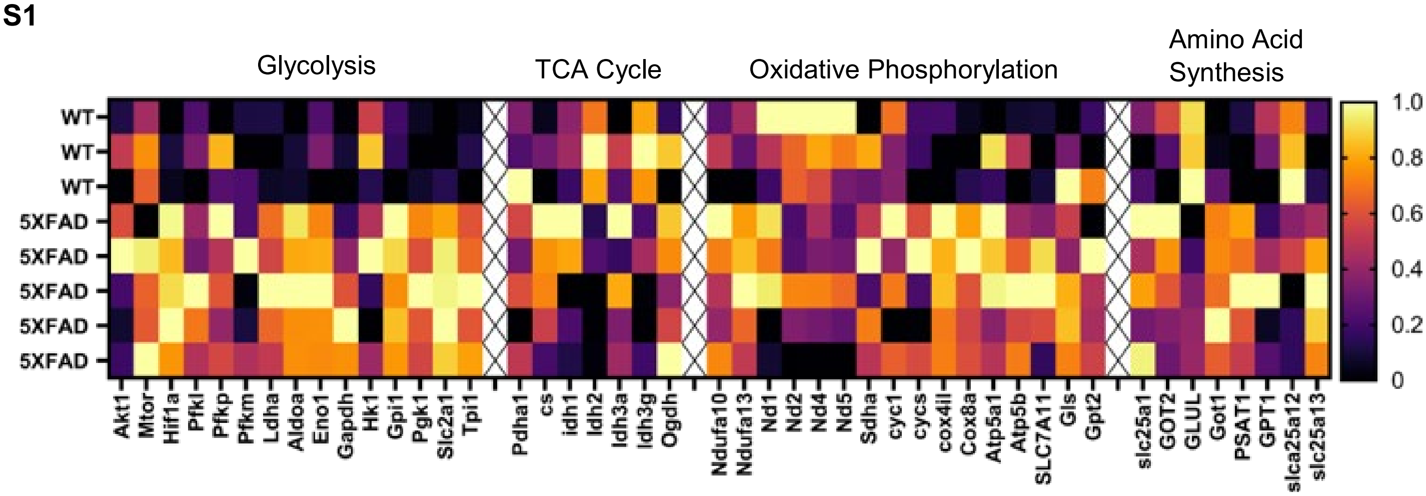


Figure S1. Microglia metabolism genes differentially expressed between 5XFAD and WT mice. Differential expression of metabolism-related genes of previously published microarray data (GSE65067). The wildtype and 5XFAD subsets were utilized for comparison using GEO2R. Differentially expressed genes relating to metabolic pathways (glycolysis, TCA cycle, oxidative phosphorylation, and the electron transport chain) were targeted and are presented in the heatmap. Each gene is normalized to the highest expression value within the comparison.

**S2**

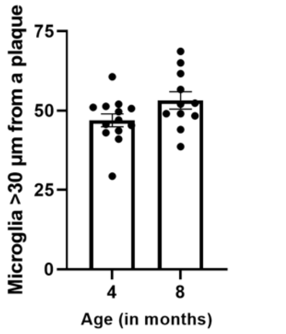


Figure S2. No difference in the number of non-plaque associated microglia. From the cluster analysis of microglia around a plaque (see images in Figure 2A). The number of microglia greater than 30 µm from a plaque was quantified. Data is presented ± SEM and each point represents 3 images. Unpaired t-test, p>0.05.

**S3**

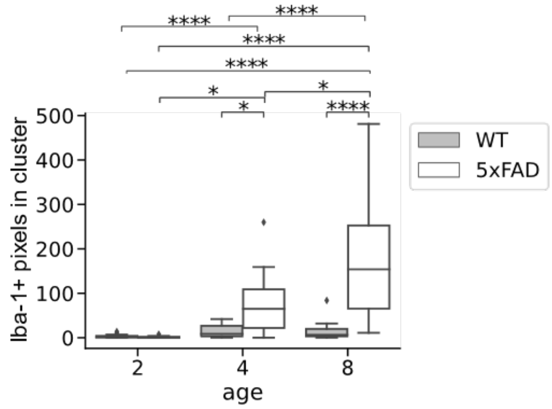


Figure S3. Quantification of the Iba-1$^{+}$ pixel count that contains the third component by age and genotype (C). ANOVA with Tukey comparisons; ****$p<0.0001$, ***$p<0.001$, **$p<0.01$, *$p<0.05$.
